# Analysis of behaviour in the Active Allothetic Place Avoidance task based on cluster analysis of the rat movement motifs

**DOI:** 10.1101/157859

**Authors:** Tiago V. Gehring, Malgorzata J. Wesierska, Daniel K. Wójcik, Eleni Vasilaki

## Abstract

The Active Allothetic Place Avoidance test (AAPA) is a useful tool to study spatial memory in a dynamic world. In this task a rat, freely moving on a rotating circular arena, has to avoid a sector where shocks are presented. The standard analysis of memory performance in the AAPA task relies on evaluating individual performance measures. Here we present a new method of analysis for the AAPA test that focuses on the movement paths of the animals and utilizes a clustering algorithm to automatically extract the stereotypical types of behaviour as reflected in the recorded paths. We apply the method to experiments that study the effect of silver nanoparticles (AgNPs) on the reference memory and identify six major classes of movement motifs not previously described in AAPA tests. The method allows us to analyse the data with no prior expectations about the motion to be seen in the experiments.

## Introduction

Navigation in a stable environment is based on allothetic or idiothetic memory, or both. However, these two kinds of memory are brought into a conflict when relevant and irrelevant (misleading) information are presented simultaneously (***Bures et al., 1997***; ***Fenton et al., 1998***). Formation of proper allothetic memory in such conditions requires segregation of information that involves cognitive coordination processes (***Wesierska et al., 2005***; ***Phillips and Silverstein, 2003***).

The Active Allothetic Place Avoidance (AAPA) test, also known as the Carousel maze test (***Stuchlík et al., 2013***; ***Dockery and Wesierska, 2010***; ***Cimadevilla et al., 2001***; ***Stuchlik et al., 2014***) is an experimental setup to study formation of allothetic memory in the presence of conflicting information. In this test animals are placed on a dry circular arena where they can freely walk. In such conditions they have to learn to avoid a shock sector, which is not marked physically but is fixed with regards to the distal, relevant cues from the room, in the presence of misleading proximal cues from the arena. The entrance to this sector is signalled by an application of a short lasting mild electric shock to the rat’s paws, which is repeated at short time intervals until they leave this sector. Thus, proper navigation in the AAPA task, requires on-going active segregation of the irrelevant local cues (e.g. faeces, urine) from the arena and use of only the distal relevant cues from the room.

The AAPA task is a variation of the passive Place Avoidance Task (***Bammer, 1982***; ***Haroutunianet al., 1985***). In this task animals are placed in a chamber divided into two compartments, dark and light. Here, they also need to avoid shocks, which are presented in the dark compartment, but contrary to the AAPA, they do so by suppressing their activity and remaining in the light compartment.

The performance in the AAPA task has been shown to be strongly hippocampal dependent (***Cimadevilla et al., 2000***) and more sensitive to its unilateral blockade (***Cimadevilla et al., 2001***; ***Wesierska et al., 2005***), than the performance in the Morris Water Maze (MWM) (***Vorheesand Williams, 2006***; ***Morrisetal., 1982***), which is a commonly used navigation task in which animals are placed in a pool of water and have to find a hidden platform to escape the water. The difference may follow from the fact that in the MWM only distal cues are available and useful for animals to orient themselves (***Morris, 1981***). Another advantage of the AAPA task, compared to the MWM, is that swimming is less natural for the rat than freely moving on the stable ground of the arena.

Commonly used measures to asses memory in the active place avoidance task (***Stuchlik et al., 2007***; ***Stuchlik and Vales, 2008***; ***Wesierska et al., 2009***, ***2013***) are: the total number of entrances to the shock sector, the number of shocks received, the time to the first shock, and the longest time of shock avoidance (the total path length and linearity of the path are considered here as measures of locomotor activity, not memory). Although very useful, these performance measures of memory do not give a direct indication about how the animals behave during acquisition of memory and how their behaviour changes during a session. In the case of spatial memory testing in the Morris Water Maze the limitation of single performance measures has been identified long time ago (***Gallagher et al., 1993***; ***Dalm et al., 2000***). The individual measures alone, like time or distance to the platform, simply cannot account for the variety of different behaviours or strategies observed in the experiments. Therefore, other analysis methods based on the classification of the swimming paths of the animals have been proposed over the years (***Wolfer and Lipp, 2000***; ***Graziano et al., 2003***). These methods combine a number of different measures of the trajectories to define a set of classes of behaviour. In ***Graziano et al.*** (***2003***) an automated classification method for MWM trajectories is also presented. Their method is based on a supervised machine learning algorithm which has to be first trained using manually labelled data, but which can then be used to classify other datasets. However, these classification methods are typically focused on assigning one trajectory to a single class of behaviour, which cannot always be reliably done because animals frequently change the patterns of their movement within trials. This makes it difficult to unambiguously map one trajectory to a single type of behaviour.

In ***Gehring et al. (2015)*** a more granular method for classifying the MWM trajectories was presented. In that study multiple overlapping segments of the swimming paths, instead of the complete paths, were classified. This made it possible to identify changes of exploration strategy within a single trial and to highlight subtle behavioural differences between groups of tested animals where other methods failed. The classification of the swimming path segments was done in a semi-automated fashion: a partial set of the MWM data was manually classified and used to constrain a clustering algorithm and determine to which classes the clusters belong. The method developed there falls therefore to the class of semi-supervised algorithms.

Here we show how the classification method developed by (***Gehring et al., 2015***) can be generalised to other experimental setups and how it can be turned into a completely unsupervised method, i.e., the classification is done based only on structural similarities of the trajectories. Using the original record of the rats moving in the AAPA test as the specific example we develop an analysis method for the AAPA experiments that is complementary to standard performance measures and which can give further insight into how the behaviour of animals changes over time and differs between groups of animals. As a case study and in order to validate the method, a set of the AAPA experiments investigating how silver nanoparticles (AgNPs) affect the spatial memory of rats is analysed here with the proposed method and compared against the standard approach. In these experiments it was found that the rats administered with silver nanoparticles, unlike the non-treated control rats, presented memory impairment (in preparation).

The method developed here, as in the case of the MWM swimming paths, is based on analysing trajectory segments instead of the full trajectories. Trajectories are therefore first split into segments which are grouped into different clusters with the help of a clustering algorithm. However, contrary to the analysis method for the MWM trajectories, the method developed here does not require any labelled data or manual data classification of any sort. Behavioural classes of interest, each one of them mapped to one cluster, do not have to be selected in advance but are instead identified by the clustering algorithm itself. These differences turn the method into an unsupervised algorithm; they are significant because they allow to analyse the data both faster, since no manual labelling is necessary, and without any previous knowledge about the types of behaviour seen in the experiments. The proposed analysis method is first introduced and then applied to a data set from AAPA experiments in order to demonstrate that it can be successfully used to identify different types of behaviour. The observed differences in behaviour between treated and untreated animals are then compared with standard analysis results. It is shown that both give consistent results but that our approach provides complementary information not apparent from the individual measurements alone.

In what follows our proposed analysis method is first introduced and then applied to a data set in order to demonstrate that it can successfully identify stereotypical types of behaviour in the data. The observed differences in behaviour between treated and untreated animals are then compared with standard analysis results to make sure that the results are consistent. This is followed by a discussion of the results and future work perspectives. Finally, a detailed description of our proposed method is presented.

## Results

We propose a new analysis method for the active place avoidance experiments which focuses on identifying stereotypical behavioural patterns of animals. Our method is based on segmenting the trajectories of the animals and then grouping similar segments by features such as their position in relation to the sector where shocks were applied, geometry, and movement speed, among others.

As a case study the method is applied to a set of experimental data acquired at the Nencki Institute of Experimental Biology, Warsaw. In the experiments 20 rats were submitted to the AAPA task and their trajectories were recorded. Of those animals 10 were administered orally with silver nanoparticles (AgNPs); the other 10, which received water, were the untreated control group. Each animal was submitted to 5 sessions of 20 minutes each during which they could move freely in a circular rotating arena and had to learn to avoid a shock sector which was fixed with respect to the room. This was followed by one 20 min test trial in which the shock sector was deactivated. Effective performance of place avoidance indicate proper spatial memory functioning.

Previous performance analysis methods of the AAPA experiments typically rely on comparisons of individual performance measures, which can well reflect how successfully each group of animals avoided the shock sector, but offer little insight into the differences in their behaviour. The method we introduce here, on the other hand, is able to identify and highlight the differences in behavioural patterns of the studied animals. This not only leads to a better understanding of how the animals learn to navigate within the arena, but also makes it possible to identify differences in behaviour between the different treated groups as well as during memory acquisition and retrieval, which would otherwise not be possible.

### Standard performance measures

Standard performance measures for the AAPA usually compare the number of entrances to the shock sector, number of shocks received by an animal during a session, the time to the first entrance, and the longest avoidance time (see ***Wesierska et al.*** (***2013***) for a detailed description of these and other statistics). Figure 1 shows the above performance measures for rats treated orally with silver nanoparticles and the untreated control group during consecutive sessions of spatial memory acquisition (Figure 1 a-d). The results show that the treated animals made more entrances with shorter time to the first entrance and shorter maximum avoidance time within a session than the untreated ones. Contrary to differences in memory measures no difference between groups was found in speed and locomotor activity measured with the total path length during the whole session (Figure 1 e-f).

**Figure 1.**
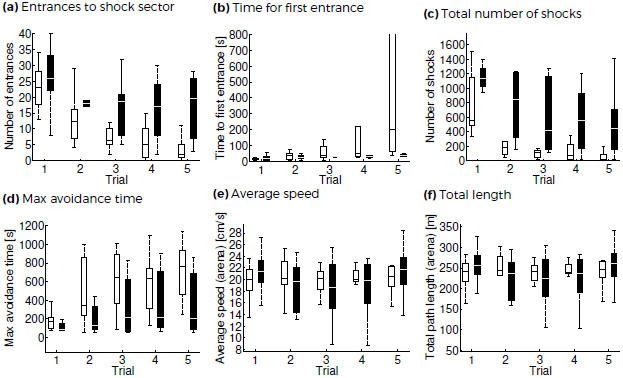
Comparison of performance between untreated control (white) and treated (black) animals over a set of 5 sessions. Boxes represent the first, second (median, shown as a band) and third quartiles; whiskers are the minimum and maximum values. (a-d): Animals in the control group are able to quickly learn how to avoid the shock sector and perform on average much better than treated animals. Average speed (e) and the total length of the trajectories (f) do not show significant differences in locomotor activity between both groups.

Although the performance measures shown in Figure 1 identify a clear difference in performance in the spatial memory task between treated and non-treated animals, they provide no indication about the types of behaviour that lead to such differences in the first place. The new analysis method we propose in what follows, on the other hand, takes a closer look at the motion of the animals and gives a better insight about how the treatment that animals were subjected to affects their behaviour.

### Classification of trajectory motifs

The recorded trajectories (120 in total) for each animal and session were segmented (Materials and Methods) resulting in 6,237 trajectory segments. A set of 11 features (Table 1) was computed for each segment. The data was then transformed using principal component analysis (PCA) and the first _*pc*_ principal components were used as new features, effectively reducing the dimensionality of the data. The data was then clustered using the MCPKmeans algorithm for different numbers of target clusters, N_*c*_. In order to select appropriate values for N_*pc*_ and N_*c*_ the clustering algorithm was run multiple times for different N_*pc*_ and N_*c*_ values and the results were then compared. The criteria for choosing N_*pc*_ and N_*c*_, summarised below, are detailed in the Materials and Methods section.

**Table 1.** Features for the data clustering of trajectory segments. For a detailed description please refer to the Materials & Methods Section.

| Feature | Unit | Reference Frame |
| --- | --- | --- |
| Angular distance to shock sector | rad | Global |
| Angular dispersion | rad | Global |
| Angular dispersion (Arena) | rad | Arena |
| Median log radius | - | Global/Arena |
| Variance log radius | - | Global/Arena |
| Trajectory centrality | % | Global/Arena |
| Median speed | cm/s | Arena |
| IQR speed | cm/s | Arena |
| Median angular speed | rad/s | Arena |
| IQR angular speed | rad/s | Arena |
| Speed change frequency | $s^{-1}$ | Arena |

1. The maximum correlation between any two clussters should not be too high (< 90% as a thumb rule). A high correlation is an indication that two or more clusters are very similar so N_*c*_ should be reduced;
2. The minimum number of elements inside a cluster is not too small: every cluster should contain at least 5 to 10% of the total number elements, although this value can be smaller when the target number of clusters is large. This is to prevent empty or close to empty clusters, which would otherwise be usually an indication that the target number of clusters or dimensions is too high.

For the purposes of our analysis we have assigned each separate cluster to a distinct behavioural class.

Figure 2 shows the maximum correlation between clusters (in %) and minimum cluster size (in % of the total number of elements) for N_*pc*_ between 5 and 7 and an increasing number of clusters, starting at N_*pc*_ = 5. As we can see, for N_*pc*_ = 5 principal components (left plot) close to empty clusters (dotted lines) and/or highly correlated clusters (≥ 90% correlation; continuous lines) always appear, which is why the results were discarded. The results for N_*pc*_ = 6 (right plot), on the other hand, show both a more homogeneous distribution of cluster sizes and a smaller maximum correlation between the clusters for N_*c*_ = 5 or 6. The results for N_*pc*_ = 7 show also similar results, albeit only for 5 target number of clusters. Because of this fact and because having a smaller number of dimensions is always desirable to avoid over-fitting, N_*pc*_ = 6 was adopted here. For the target number of clusters N_*c*_ = 6 was adopted since there is no appreciable increase in the maximum correlation between clusters between N_*c*_ = 5 and N_*c*_ = 6, which suggests that N_*c*_ = 5 contains one cluster that can be separated into two and that N_*c*_ = 6 is the more natural choice.

**Figure 2.**
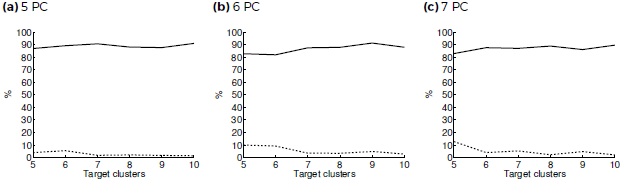
Clustering results for different number of principal components and target number of clusters. Continuous lines: maximum correlation between clusters; Dotted lines: minimum cluster size (in % of the total number of trajectory segments). Plots A-C show the results for 5 to 7 principal components.

Figure 3 shows two example segments of trajectories for each of the resulting 6 clusters. Segments are shown both in the room coordinates (as registered by the camera; left circle) and the rotating arena reference frame (real path swept by the animals; right circle); the movement speed in the arena reference frame is also shown for each one of the segments. A description of the observed behavioural traits of each cluster is given below. Table 2 gives also some statistics for each cluster, such as the relative size and percentage of segments that start or end up within the shock sector.

**Table 2.** Clustering statistics showing relative number of elements (in % of the total number of elements) and percentage of elements beginning or ending at the shock sector. Segments beginning at the shock sector are associated with types of behaviour after the animal received one or more shocks. Segments ending in the shock sector are related to behaviour just before the animal (most likely) receives a shock.

| Cluster | Elements | Entering shock | Leaving shock |
| --- | --- | --- | --- |
| 1 | 16.8% | 0.2% | 3.5% |
| 2 | 16.5% | 85.1% | 37.6% |
| 3 | 10.9% | 3.8% | 36.6% |
| 4 | 29.2% | 0.1% | 0.1% |
| 5 | 12.2% | 64.0 % | 31.9% |
| 6 | 14.4% | 1.7% | 5.8% |

**Figure 3.**
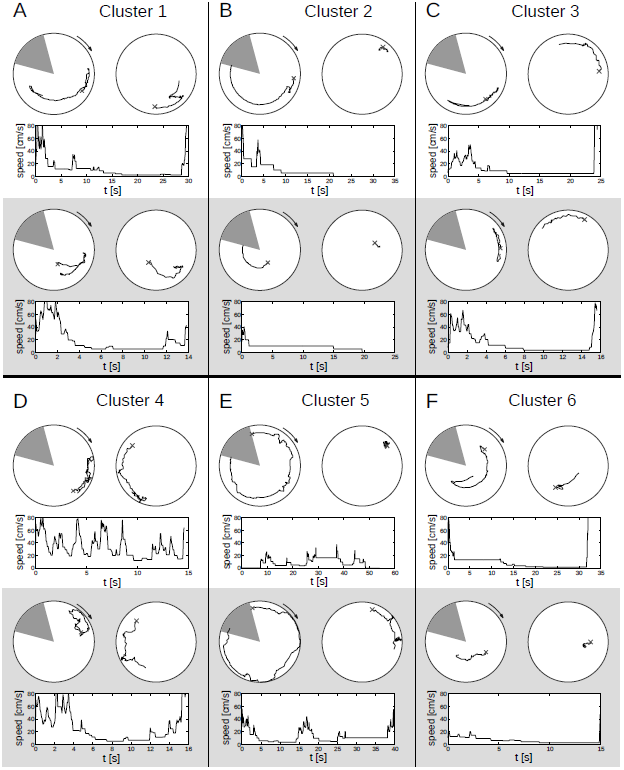
Example trajectory segments for the six resulting clusters. Two examples are shown for each cluster (white and shaded regions). Top left figures show the trajectories in the room reference frame. Top right plots show the trajectories compensating for the rotation of the arena (arena reference frame). Lower plots show the speed of the rats in the arena reference frame. The shaded triangular region marks the shock sector and crosses — the starting positions of the animals.

### Classes of behaviour

Here we describe briefly the behaviour associated with each of the resulting clusters. These classes of behaviour were not predefined but rather identified automatically by the clustering algorithm. The descriptions for each class are based solely on the observed traits of each of the computed clusters.

#### Class 1: movement inwards opposite to the shock sector

Animals move to more central points in the arena and try to stay on the opposite side of the shock sector.

#### Class 2: passive until shock sector

Animals move very little around the arena and sit mostly at one position, ending frequently in the shock sector (Table 2).

#### Class 3: movement opposite to the shock sector

Paths focused on the extreme opposite of the shock sector.

#### Class 4: random movement opposite to the shock sector

Relatively chaotic paths concentrated on the right/upper half of the arena, on the right side and immediate vicinity of the shock sector.

#### Class 5: passive until shock sector, longer paths

Similar to class 2 but producing longer paths. Animals sit mostly in one position, frequently completing a full revolution that starts and ends at the shock area (Table 2).

#### Class 6: movement inwards and to the centre of the arena

Animals move inwards and explore the more central parts of the arena.

Classes 1 and 3 are clearly the most efficient since animals actively try to stay away from the shock area. Conversely, classes 2 and 5 are the most inefficient because animals mostly stay still and never try to avoid running into the shock sector.

### Comparison of treated and control groups

Figure 4 shows the distributions of classes of trajectory segments for treated and non-treated animals. For every animal in a given group the trajectory was segmented (Materials and Methods) and the segments were attributed to one of the six different classes of motion by the clustering process. This gave an estimate of the percentage of the time that the animal adopted each of the classes of behaviour for the session. These values were then averaged for all 10 animals in each group. The differences between the groups were then checked for significance using the Friedman test. Resulting p-values for the Friedman test are shown in the plots for each cluster.

**Figure 4.**
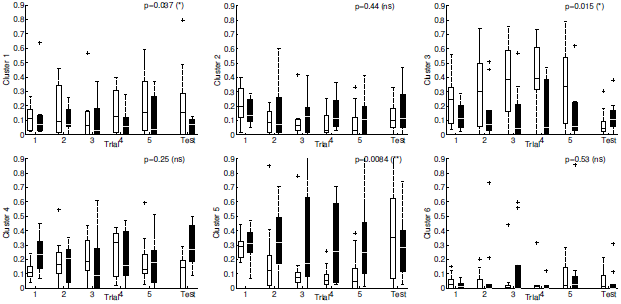
Distribution of segments from each cluster for control (white boxes) and treated (black boxes) animals for 5 days of spatial memory acquisition (with active shocks) and 1 test session on memory retrieval (without shocks) in the AAPA task. Boxes represent the first and third quartiles of the data, lines the median, crosses the outliers, and whiskers the minimum and maximum values. The Friedman test (Materials and Methods) was used to compare both groups of animals over all sessions; p-values are shown on the top right.

For 3 of the 6 clusters significant differences (p ≤ 0.05) were found between the control and silver-nanoparticles treated groups. Animals in the treated group, contrary to control, prefer to stand still on the arena and in many cases their trajectories ended in the shock sector (classes 5 and 2).

Animals in the untreated control group, significantly more often than treated rats demonstrated a behavioural pattern in which animals sit still for a while but then move away from the shock sector as they approach it (classes 1 and 3). This suggests that these animals are aware of the location of the shock sector and adopt a strategy in which they have to move around as little as possible.

The properties of the behavioural patterns obtained with cluster analysis show that treated animals significantly more often ended their motion in the shock sector than the control untreated rats. This is due to them adopting the less efficient strategy of class 5 more frequently, and exhibiting the efficient strategies of classes 1 and 3 less. It confirms the results obtained with standard measures for AAPA that treated, contrary to untreated animals presented spatial memory impairment.

## Discussion

We have presented a new analysis method for the AAPA test experiments. The method relies on splitting the recorded trajectories of the animals in the arena, computing a set of features for each resulting trajectory segment, and then using a clustering algorithm to identify similar behavioural patterns in the data. This is a generalisation of the method devised previously ***Gehring et al.*** (***2015***) for the analysis of the Morris Water Maze experiments. There, however, the classes of behaviour were predefined and a partial set of pre-labelled data was used to classify the trajectories. This effectively led to a semi-automated or semi-supervised classification method. The method presented here, on the other hand, offers a completely unsupervised approach to the classification; behavioural classes of interest do not have to be defined beforehand and no manual labelling of data is necessary.

As a case study the new method presented here was applied to a data set consisting of trajectories of 20 animals recorded in the active place avoidance test on allothetic spatial memory function. Half of the animals were treated with silver nanoparticles, the other half was the untreated control group. Although standard memory measures described earlier show an impairment of spatial memory in treated animals, the difference in performance between the two groups becomes much more evident when their behavioural patterns are compared. Impairment of avoidance in treated animals shows as poor recognition of the position of the shock sector in the room frame coordinates and diminished ability of learning efficient strategies to avoid it. Treated animals show instead a higher tendency for sitting in one position in the arena until entering the shock sector. The new method identified six types of distinct behaviours which, to the best of our knowledge, were never described in the literature before.

This work shows how machine learning algorithms can be applied to behavioural data to find patterns and create a method for identifying stereotypical navigation strategies. The semi-supervised method developed previously for the MWM was extended to a fully unsupervised context and generalised to work with another completely different experimental setup. This shows that the method is general enough and that it can be extended to further types of experiments. Although in the AAPA test, unlike the MWM test, the trajectory has usually been considered a measure of free locomotor activity, not memory, here we show that one can derive useful measures of memory from the trajectory. The software tools implementing the present analysis, which are based on the tools developed previously for the MWM, are freely available and can be used as a basis for creating more sophisticated tools or to generalise the method to other behavioural setups. We note that the circular shape of the arena is incidental to the analysis. One could apply the approach presented here to regions of arbitrary shapes, indeed, the segmentation of trajectories into pieces of equal length makes this method amenable even to open field studies or fields of classical square shapes.

One possible point of criticism of the method presented here is that the classification method depends on relatively fine tuned features which have to be defined for each type of experiment. In order to apply the method to other experimental setups, appropriate features that measure relevant geometrical aspects of the trajectories or their relative position relative to an objective (such as the escape platform in the MWM or a special area, e.g. a sector to be avoided in the place avoidance task) have to be defined. This problem is minimised to some degree here by using a larger pool of features and a feature extraction method in the form of PCA. This is done to reduce the dimensionality of the data set and to find a reduced set of most relevant features that can account for most of the data variability. Nevertheless, the problem of first having to design appropriate features for a given experiment remains. In order to fully overcome this limitation the possibility of defining abstract measures that can be more universally applied remains to be investigated. One promising approach for achieving this is presented in ***Korz*** (***2006***), which introduces a method in which the coordinates of the paths of animals in the MWM are used to define a new set of features. In his work PCA is also used to reduce the resulting high dimensional feature space and it is shown that the first few principal components are enough to account for most of the variability in the data. The same approach was not adopted here since trajectories were classified not only by their geometrical aspect but also by other factors such as their relative position in the maze and the movement speed of the animals. However, a more general approach that, for example, combines automatically defined features from geometrical aspects with hand tuned positional measures could be investigated in the future.

## Materials and methods

### Experimental setup

The experiments using the Active Place Avoidance Test were conducted at the Nencki Institute of Experimental Biology, Warsaw, Poland. The same basic experimental setup as described in ***Wesierska et al.*** (***2009***) was used. The setup consisted of an aluminium circular arena 80 cm in diameter and a 2 cm rim which rotated with one revolution per minute. The arena was positioned 80 cm over the floor and placed in the centre of a 3×4 meter lightly lit room which contained many stable external visual cues. Infrared light-emitting diodes (LED) for tracking the position of the animals and a 25G (0.50 mm) hypodermic needle electrode were attached to the backs of the rats. A second LED was attached to the periphery of the arena. It allowed monitoring the position of the rat by the infrared TV camera which was connected to a computer system. The experimental setup is shown schematically in Figure 5.

**Figure 5.**
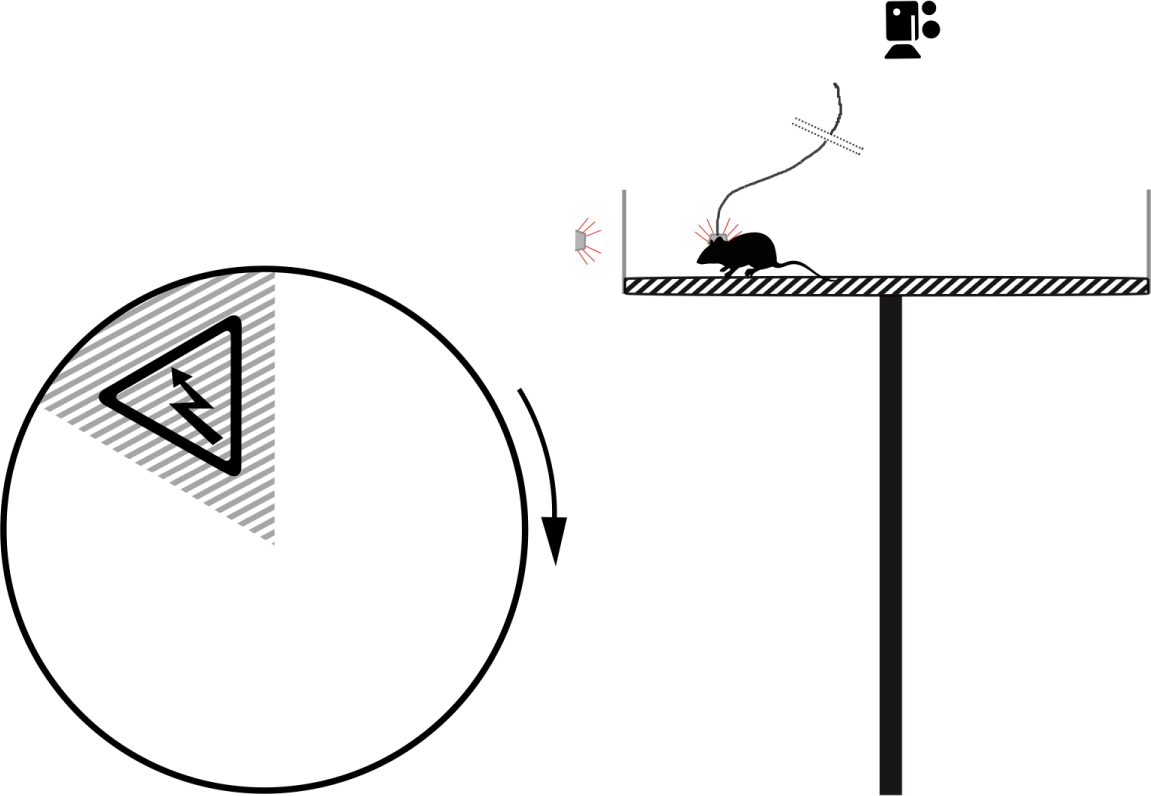
The Active Place Avoidance setup. Animals are placed on top of an elevated arena which is slowly rotating (1 revolution per minute). They can move freely around the arena but need to learn to avoid the shocks, which are delivered on sector, which is fixed according to the distal room cues. If they enter sector to be avoided, a short lasting low current pulse is delivered to their paws and repeated with a delay until they leave this sector. The position of the animals is tracked with LEDs 1 and 2, and a top-mounted camera.

Five recording sessions of 20 min each with a fixed shock sector (in the room coordinates) were performed over a set of five consecutive days. This was followed by a test trial five days later where the shock sector was not active. For the trials with an active shock sector animals received a short (0.5 s) constant current pulse whenever they entered a predefined 6° shock sector, which remained fixed across the trials. The amplitude of the shock pulses varied between 0.2 and 0.5 mA and was determined individually for each animal so that shocks did not make the animal freeze or induce attempts to escape the arena. Shocks were repeated every 1.5 s until the animal left the shock sector. The position of the animal and the current state of the electrode (shock active or not) was recorded at 25 Hz using commercial software (Bio-Signal Group, New York).

### Animals and treatment

Twenty naïve adult (2.5 month-old) male Wistar rats, weighing 270–310g, were obtained from the breeding colony of The Center of Experimental Medicine of the Medical University of Bialystok, Poland. They were accommodated in transparent plastic home cages, four animals per cage, under standard conditions (a constant temperature of 22° C, 12:12 light/dark cycle, humidity at 50%). Water and food were available in the cages ad-libitum. 28 days before the experiments ten of the animals were treated orally with silver nanoparticles (experimental group) and ten with water (untreated control group).

All the manipulations were done according to the European Community directive for the ethical use of experimental animals and the Polish Communities Council for the care and use of laboratory animals.

### Data analysis

The recorded trajectories of the animals were exported from the data acquisition system as text files and further processed by custom data analysis software written in Matlab. The data analysis method employed here is an extension and generalization of the method described in ***Gehring et al.*** (***2015***). The classification used in that work was based on a semi-supervised clustering algorithm which made use of a partial set of labelled data to constrain the clustering algorithm and to map clusters to one of the predefined classes of behaviour.

In the present work, data analysis was also based on a classification of trajectory segments into different stereotypical types of behaviour, however, classes of behaviour were not predefined. Also, no labelling of data of any kind was performed, that is, the classification performed here was completely unsupervised.

The analysis done here consisted of the following steps:

1. splitting the trajectories of the animals into shorter segments;
2. computing a set of features for each segment and reducing the dimensionality of the data;
3. clustering the data;
4. analysing the distribution of the resulting clusters for both groups of animals;

#### Segmentation of trajectories

The main focus of our analysis was to understand how the strategies of animals for avoiding the shock sector evolve over time (between sessions) and differ between treated and untreated animals. Therefore, in the first step the recorded trajectories were split into fragments delimited by entrances/exits from the shock sector. That is, only the parts of the trajectories not falling in the shock sector were considered. Since the length of the trajectories between shocks varied widely, from the order of a few seconds up to the duration of the trial (20 min), and since the animals during this long time usually display multiple types of behaviour, these fragments were split further.

The objective of the second segmentation step was to isolate the different behaviours found in a trajectory and to generate a more uniform distribution of segment lengths, to facilitate classification. The second segmentation used changes in the angular speed as criteria for splitting the trajectory segments. This is because in the Active Allothetic Place Avoidance Test animals have to move in the angular direction in order to evade the shock sector. Therefore, changes in the sign and magnitude of the angular speed were taken as the delimiting points of the segments. More formally, in the second segmentation step trajectory points were processed sequentially and added to a sub-segment until the difference between the local and median angular speed of the sub-segment (recomputed for each new added point) exceeded 0.6 rad/s. Segments shorter than 5 seconds were discarded in further analysis.

The two segmentation steps are shown schematically in Figure 6. From the original 120 trajectories, 1,741 segments were generated after the first segmentation step and 6,237 after the second. Other statistics of the two segmentation steps can be seen in Table 3.

**Table 3.**
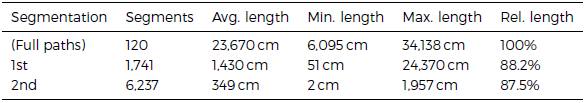
Segmentation of trajectories statistics. All lengths are measured in the arena (rotating) reference frame. The last column shows the total length of the resulting segments compared to the input, i.e., without the short segments that were discarded.

| Segmentation | Segments | Avg. length | Min. length | Max. length | Rel. length |
| --- | --- | --- | --- | --- | --- |
| (Full paths) | 120 | 23,670 cm | 6,095 cm | 34,138 cm | 100% |
| 1st | 1,741 | 1,430 cm | 51 cm | 24,370 cm | 88.2% |
| 2nd | 6,237 | 349 cm | 2 cm | 1,957 cm | 87.5% |

**Figure 6.**
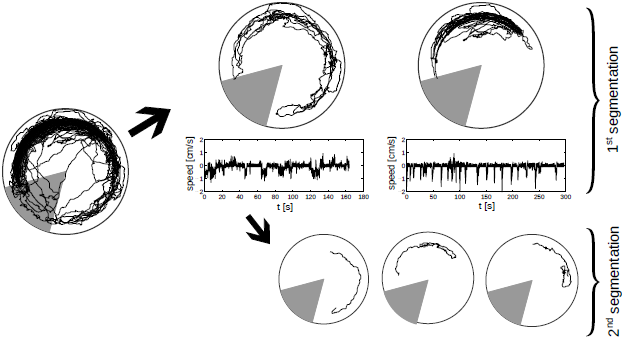
Two-step segmentation of the trajectories. In the first step trajectories are split into segments containing only the parts of the paths not falling inside the shock sector. In the second step, sudden changes in the angular speed (middle plots) are taken as the delimiting points of the segments.

### Computation of features

For each trajectory segment a set of 11 features was computed. The resulting data space was then reduced to a smaller dimensional space using Principal Component Analysis (PCA). The resulting data was then fed to a clustering algorithm.

This section describes the 11 features, that measure different geometrical and positional aspects of the segments, used in the classification (Table 1). Some features are computed using the room reference frame, that is, the coordinates including the rotation of the arena; other features are computed in the rotating reference frame, i.e., using the real paths swept by the animals on the arena. A detailed description of each feature is given in what follows.

#### Angular distance to shock sector

This value measures the angular distance from the centre of the shock sector in the room coordinate frame to the angular centre of the segment. The latter is computed by adding the position vectors of each sample in the trajectory (i.e. the vector to the centre of the arena) and then taking the angle of the resulting vector relative to the middle shock sector angle. If the resulting angle is negative, 2π is added to it so that the resulting values are in the [0,2π) range.

#### Angular dispersion

The angular dispersion measures the angular spread of the trajectories in the room coordinate frame. It is here defined as the differences between the maximum and minimum angles of the position vectors of each data sample in the trajectory.

#### Median/IQR of the log-radius

These values are calculated from the trajectory by computing the distance to the centre of the arena for each data sample, taking the logarithm and then computing the median (interquartile range) of the values. The median and IQR were chosen over the mean and standard deviation because they are less susceptible to outliers.

#### Trajectory centrality

Measures the relative amount of time that the animal spends at the more central regions of the arena. The value is computed by computing the length of the trajectory falling within a concentric circle with a radius of 75 % of the radius of the arena and dividing this value by the total length of the trajectory.

#### Median/IQR speed

The speed at each trajectory point is computed and the median/interquartile-range of the resulting values is then calculated.

#### Median/IQR angular speed

The angular speed (relative to the centre of the arena) at each trajectory point is computed and the median/interquartile-range of the resulting values is then calculated.

#### Speed change frequency

Measures the number of times that the speed changes abruptly within the segment. Calculated by counting the number of times that the (absolute) speed crosses25 % of the median speed of the segment.

The 11 features computed for each trajectory segment were not used directly for clustering the data. This is because in a high dimensional vector space the distance between elements tends to be very similar, making it difficult to find meaningful clusters (***Aggarwal et al., 2001***). In order to overcome this problem without having to explicitly select a small subset of features, Principal Component Analysis (PCA) (***Wold et al., 1987***) was used. PCA transforms N possibly correlated sets of features into N linearly uncorrelated variables, or principal components. The principal components are defined so that each successive component points to the direction that maximizes the variance of the data. That is, the first principal component accounts for most of the variability of the data, followed by the second, and so on. Therefore, keeping the first few principal components and projecting the old feature values onto them one can reduce dimensionality of the data.

Here the data were clustered multiple times using different numbers of principal components. The criteria used to select the appropriate number of components, or the dimension of the resulting feature space, are described below.

### Clustering

We used Metric Pairwise Constrained K-Means (MPCKMeans) clustering algorithm (***Bilenko et al., 2004***). It is based on the classic K-Means algorithm (***MacQueen, 1967***; ***Hartigan and Wong, 1979***) but supports features such as metric-learning and constrained clustering, which technically makes it a semi-supervised algorithm. The latter feature was not used here, but constraints can be very useful if, for example, a set of predefined classes of behaviour and prelabelled data is available (see for example ***Gehring et al.*** (***2015***) where this feature was used). Since no labelled data was used here our method is in practice completely unsupervised.

Although it would have been possible to use a standard K-Means algorithm for the analysis here, the MPCK-means was chosen instead in order to maintain consistency with the previous work on the MWM trajectories (***Gehring et al., 2015***) and because of its support for some additional features, such as clusters of different shapes, made possible by a cluster-dependent metric function updated at each cluster iteration. This is in contrast to the standard K-means algorithm which uses a single metric function which usually leads to a more homogeneous distribution of cluster sizes and shapes. Another important property of the MPCK-means algorithm which makes results easier to compare and recall is its deterministic nature. In classic K-means results are usually non-deterministic because of the use of random initial conditions.

### Number of principal components and target number of clusters

One of the difficulties in using a clustering algorithm is choosing the appropriate target number of clusters. To choose appropriate target number of clusters and the number of principal components (or features) for clustering the data the following criteria were used:

1. The maximum correlation between any two clusters should not be too large (≤ 90% as a thumb rule). A large correlation between clusters means that two or more clusters are too similar and therefore redundant;
2. The minimum number of elements inside a cluster is not to small (≥5 - 10% of the total elements, although the value depends on the number of target clusters, or classes). This is to avoid having clusters that are empty or close to empty;

The correlation between two clusters was computed by averaging the correlation between N elements closest to the centroids of each cluster, where N is the size of the smallest cluster. The data was clustered using different number of principal components and number of clusters. The results that fulfilled both of the above conditions with the minimum number of principal components (or dimensions) and maximum number of clusters (or types of behaviour) were then adopted.

#### Statistics

Multi-factor testing of variance was done using a Friedman test (***Siegel, 1956***), a nonparametric test that is well suited for data that is not normally distributed. The Friedman test is also a matched test, and can control for experimental variability among subjects. In our case the same animals were analysed over multiple sessions, which show a gradual change in behaviour over time. The variability between sessions, that affects all animals, was not taken into account. The p-values shown in our analyses answer the question: if the effect of different treatments (untreated control vs. silver-nanoparticles treatment) is identical, what is the chance that a random sampling would result in the distribution of values as far apart as observed? Small p-values (< 0.05 in our analyses) lead us to discard the null hypotheses that the results are identical and differences are only due to random sampling.

